# A New Perspective on Individual Reliability beyond Group Effects for Event-related Potentials: A Multisensory Investigation and Computational Modeling

**DOI:** 10.1101/2021.09.06.459195

**Authors:** Zhenxing Hu, Zhiguo Zhang, Zhen Liang, Li Zhang, Linling Li, Gan Huang

## Abstract

The dominant approach in investigating the individual reliability for event-related potentials (ERPs) is to extract peak-related features at electrodes showing the strongest group effects. Such a peak-based approach implicitly assumes ERP components showing a stronger group effect are also more reliable, but this assumption has not been substantially validated and few studies have investigated the reliability of ERPs beyond peaks. In this study, we performed a rigorous evaluation of the test-retest reliability of ERPs collected in a multisensory and cognitive experiment from 82 healthy adolescents, each having two sessions. By comparing group effects and individual reliability, we found that a stronger group-level response in ERPs did not guarantee a higher reliability. Further, by simulating ERPs with a computational model, we found that the consistency between group-level ERP responses and individual reliability was modulated by inter-subject latency jitter and inter-trial variability. The current findings suggest that the conventional peak-based approach may underestimate the individual reliability in ERPs. Hence, a comprehensive evaluation of the reliability of ERP measurements should be considered in individual-level neurophysiological trait evaluation and psychiatric disorder diagnosis.

## 1. Introduction

Event-related potentials (ERPs) are noninvasive electrophysiological measures of indexing a range of sensory, cognitive, and motor processes involved in human brain activity. In clinical and translational applications of ERPs, a key challenge is to identify a reliable and valid mapping between individuals’ brain activation and their perceptual or cognitive capacities (Nelson and Guyer, 2012). Measurement reliability is the prerequisite for clinical applications of ERPs, such as assessments of meditative practice using sensory-evoked potentials (Cahn and Polich, 2006) or diagnoses of psychiatric cognitive dysfunction by cognitive ERPs like P300 (Polich, 2004), and studies concerning reliability have received more attention recently (Dubois and Adolphs, 2016; Höller *et al*., 2017; Noble *et al*., 2019; Croce *et al*., 2020).

Originating from the field of psychometrics, reliability reflects the “trustworthiness” of a measure and denotes the extent to which a measure will yield a reproducible difference between individuals (Kraemer, 2014). The importance of reliability in the research of individual difference cannot be overstated, regardless of the data analytics approaches used (e.g., correlational analysis or machine learning). In correlational analysis, the ability to find correlations between brain activation and cognitive behavior depends on the reliability of these measures (Goodhew and Edwards, 2019). In other words, the maximum possible correlation is constrained by the reliability of the individual measures used to calculate the correlation (Spearman, 1910). In machine learning-based individualized prediction, reliability has been proved mathematically to provide a lower bound on predictive accuracy (Bridgeford *et al*., 2020).

Since the first systematic study on the reliability of ERPs (Segalowitz and Barnes, 1993), numerous studies have evaluated the test-retest reliability of ERP amplitude and the latency elicited from a variety of experimental paradigms (Cassidy *et al*., 2012; Cruse *et al*., 2014), but the primary focus has always been restricted to narrow time windows around ERP peaks (Thigpen *et al*., 2017; Cruse *et al*., 2014; Ip *et al*., 2018). Characteristic features, including latency, maximum amplitude, mean amplitude, and area under the window, are typically used to examine the reliability of ERPs. These ERP features are used in a machine learning model or correlation analysis to establish linkage between ERPs and cognitive/behavioral variables (Hu and Iannetti, 2019). However, such an analysis routine implicitly assumes that only the peak-related ERP measures reflect the subject-specific neurophysiological process to an external stimulus. This assumption is problematic, because the entire ERP shapes (rather than latency and amplitude of ERP peaks) are physiologically meaningful and important (Gaspar *et al*., 2011). Taking the temporal evolution of facial emotion perception as an example, the temporal shape of ERP can provide valuable clues about processing dynamics beyond what can be inferred from data restricted to ERP peaks (Van Rijsbergen and Schyns, 2009).

In fact, ERP peaks represent the strongest group effects (i.e., group-level experimental effects among different conditions/cohorts). More specifically, by comparing the ERP response with its baseline activity, or contrasting two experimental conditions (i.e., the ERP difference wave), peak-related features of well-known ERP components, like N100, N200, and P300, were claimed to be closely associated with various perceptual and cognitive variables (Sur and Sinha, 2009). Here, the focus was on significant group effects responding to one condition versus another. As a representative example relevant to this research, the P300 was found to reflect the processes involved in stimulus evaluation or categorization as evidenced by experimental manipulation; thus, it is often reasonable to ask whether peak-related features of P300 reflect an individual’s cognitive function. From this perspective, as the dominant approach in investigating individual differences in ERPs, peak-based analysis implicitly employs group-level prior information. However, from the perspective of individual difference, it remains unclear whether peak-related activity shows robustness or consistency in assessing between-subject variance(Brandmaier *et al*., 2018).

Indeed, the approach of identifying regions-of-interests (ROIs) by the strongest group effects and subsequently testing them for individual reliability was a common practice in evaluating individual differences in ERP studies, but recent studies have raised concerns that such a conventional approach may reduce the probability of detecting significant individual-level effects, especially in functional magnetic resonance imaging (fMRI) (Fröhner *et al*., 2019; Infantolino *et al*., 2018). For researchers interested in individual differences, between-subject variance in brain function is usually considered as the signal of interest rather than noise (Seghier and Price, 2018). For researchers interested in experimental effects, within-subject variance is treated as the signal of interest, and between-subject variance represents the noise that should be minimized. Those different views imply that regions eliciting greater activation (i.e., a peak at an electrode showing the strongest group-averaged activity) on group effects may not correspond to reliable individual effects, which has been thoroughly discussed in psychology recently (Hedge *et al*., 2018; Goodhew and Edwards, 2019; Fisher *et al*., 2018). To the best of our knowledge, the rationality of selecting individual difference variables based on group effects in ERP analysis has been seldom challenged. Whether and in which situation the group effects and individual reliability are consistent is still questionable. In real data, the underlying factors among different subjects are unmeasurable and cannot be adjusted at will, which makes it challenging to answer this question. Thus, a simulation model should be applied to investigate underlying factors of modulating the consistency between the group effect and individual reliability, but this investigation via computational modeling is still absent.

To address the abovementioned problems, the present study sought to examine the test-retest reliability of sensory-evoked potentials and cognitive ERPs based on the whole waveforms but not those restricted to narrow time windows around the peaks. More specifically, to test whether there is a spatial and temporal dissociation between group effects and the individual reliability result, the reliability of auditory-evoked potential (AEP), somatosensory-evoked potential (SEP), visual-evoked potential (VEP), and P300 was systematically examined by spatiotemporal decomposition and evaluation in a pointwise way (i.e., at each spatial-temporal EEG sample). Further, a dynamical system model was applied for the simulation of ERP generation to investigate the underlying mechanism explaining the real data results, in which key model parameters were varied to test their influences on the consistency between group effects and individual reliability.

## 2. Materials and Methods

### 2.1 Data collection and preprocessing

#### 2.1.1 Experimental paradigm

A total of 112 healthy subjects participated in this study, and 93 subjects (Mean_age_ = 21.1 year; SD_age_ = 2.3 years) among them attended two sessions, which were scheduled on different days, separated more than 6 days and 20 days apart on average. After removing 11 subjects whose data were corrupted with heavy artifacts, 82 subjects were included in subsequent reliability analyses. Ethical approval of the study was obtained from the Medical Ethics Committee, Health Science Center, Shenzhen University (No. 2019053). All subjects were informed of the experimental procedure, and they signed informed consent before the experiment. As illustrated in Fig. 1, the experimental paradigm was the same for the two sessions on different days. The experimental paradigm contained three types of sensory-evoked experiments (visual, auditory, and somatosensory) and a cognitive visual oddball experiment.

**Fig. 1.**
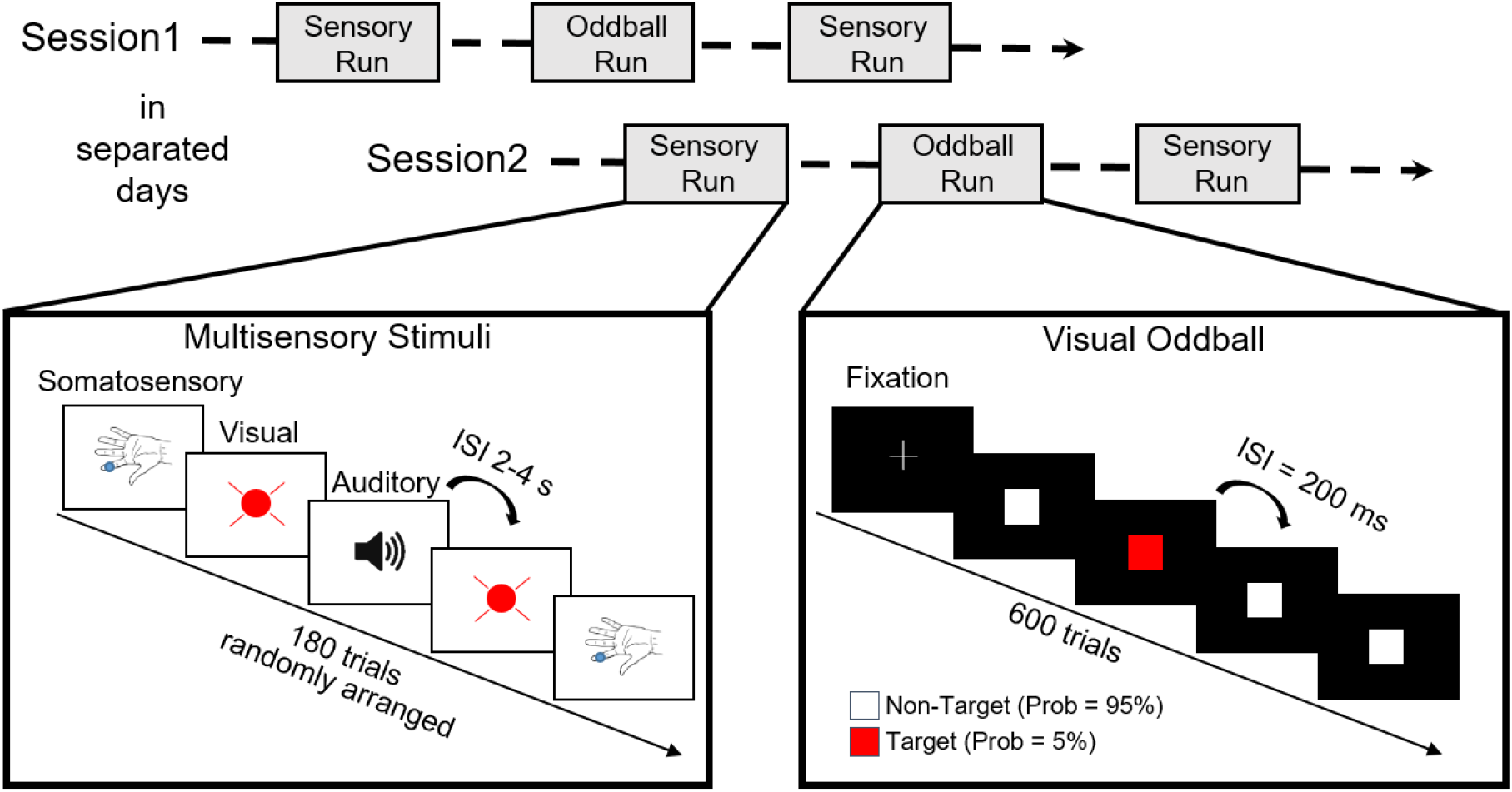
Experiment procedure. Lower Left: Sensory-evoked potentials were elicited by a random sequence of somatosensory, auditory, and visual stimuli. Auditory stimuli were brief tones produced by a speaker; visual stimuli were brief flashes produced by an LED; somatosensory stimuli were applied to the index finger of the left hand by a vibrator. Lower Right: Cognitive ERPs were elicited by the classical visual oddball paradigm with the red squares as the target stimuli and white as the nontarget stimuli on the screen.

Multiple sensory stimuli were arranged in two runs for each session. Each run consisted of 90 trials, including visual, auditory, and somatosensory vibration stimuli. These stimuli were delivered in a random order with inter-stimulus-interval (ISI) randomly distributed in the range of 2–4 s. Each stimulus lasted 50 ms. Hence, for each subject, there were a total of 180 trials of sensory stimulation in each session and 60 trials for each of the visual, auditory, and somatosensory stimuli. An Arduino Uno platform was programmed to release the three types of stimuli, which communicated with the Matlab program (The MathWorks Inc., Natick, USA) on a PC through a serial port. An light-emitting diode (LED: 3 W with light shield), a headphone (Nokia WH-102), and a vibrator (1027 mobile phone flat vibration motor) were used to generate the visual, auditory, and somatosensory stimuli, respectively.

The P300 experiment was arranged between the two runs of multiple sensory stimuli for each session. The visual oddball experiment was performed with the red squares as the target stimuli and the white squares as the nontarget stimuli on the screen. Each square lasted 80 ms, with an ISI of 200 ms. Hence, a total of 600 trials were delivered within 2 min in a run, in which the target stimuli appeared with the possibility of 5%. A subject was asked to count the number of red squares and report the result at the end of the run to keep his/her attention on the screen.

#### 2.1.2 EEG recording and preprocessing

EEG signals were recorded via a multichannel EEG system (64 Channel, Easycap) and an EEG Amplifier (BrainAmp, Brain Products GmbH, Germany). The signals were recorded at a sampling rate of 1000 Hz by 64 electrodes, placed in the standard 10-20 positions. FCz was set to be the reference. Before data acquisition, the contact impedance between the EEG electrodes and the cortex was calibrated to be lower than 20 kΩ to ensure the quality of EEG signals during the experiments.

The raw EEG data were first filtered by a 0.01–200-Hz band-pass filter and a 50-Hz notch filter. Then, bad channel interpolation was performed, and artifacts produced by eye blinks or eye movements were identified and removed by an independent component analysis (ICA) (Jung et al., 2000). For both sensory-evoked potentials (AEP, SEP, VEP) and P300, continuous EEG recordings were segmented into 1.5-second-long epochs (from −0.5 to 1.0 s relative to stimulus onset) and band-pass filtered (0.1–30 Hz). The pre-stimulus interval from −0.5 to 0 s was used for baseline correction. Grand average ERP waveforms were computed for each participant and stimulus type (visual, auditory, somatosensory, and target stimuli of the visual oddball paradigm). All EEG pre-processing steps were carried out using Letswave7 (Huang, 2019) and Matlab.

### 2.2 Reliability analysis

#### 2.2.1 Peak-based analysis and pointwise analysis

As the peak of each ERP component indicates the time point with a larger signal-to-noise ratio in the surrounding samples, peak amplitude is commonly used as a representative feature in ERP analysis. In this research, the most significant positive and negative peaks were detected by manually searching for the local maximum/minimum value in their corresponding time intervals for each subject. The mean amplitude around the peaks was not considered in this research because it is not fair to compare the reliability of pointwise analysis with the reliability of the mean amplitude, which is the average of multiple points.

Pointwise analysis was also used to examine the reliability of the ERP. More specifically, the ERP amplitude at each time point and each channel was taken as the variable for measuring individual difference. Unlike the peak-based analysis, pointwise analysis is a fully data-driven method that is performed along with the temporal and spatial domain in a point-by-point way.

#### 2.2.2 Metric of Reliability: Intraclass correlation coefficient (ICC)

ICC is a commonly used metric for reliability analysis. In this study, the reliability was measured by using ICC(A, 1) of case 2A (McGraw and Wong, 1996) to represent the absolute agreement between repeated measurements for both the peak-based and pointwise analyses for both the peak-based and pointwise analyses. The subject-by-experiment matrix was modeled by a two-way ANOVA with random subject effects (row effects), fixed session effects (column effects), and residual effects, as shown in Eq. (1), and ICC(A, 1) is calculated as Eq. (2).

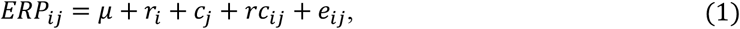

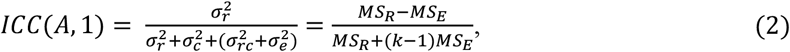

In the above two equations, *i* = 1,..,82 is used as the subscript for subjects; j = 1,2 is the subscript for multiple observations; *μ* represents the population mean for all observations; *r_i_* (row effects) is random, independent, and normally distributed, with a mean of 0 and a variance of 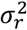; *c_j_* (column effects) is random, independent, and normally distributed, with a mean of 0 and a variance of 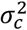; *e_ij_* (error terms) and *rc_ij_* (the interaction effects) are the residual effects with a variance of 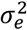 and 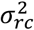. The reliability according to ICC(A, 1) was defined as the proportion of the between-subject variation over the total variation. The mean value and confidence interval of ICC(A, 1) for both peak-based and pointwise analyses were obtained 200 times by bootstrap, which involved choosing random samples with replacement from a dataset and analyzing each sample in the same way. Based on Eqs. (1) and (2), the variance in an ERP measure could be partitioned into several components as Eq. (3), which was first conceived by (Segalowitz and Barnes, 1993):

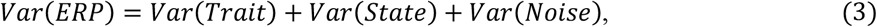

in which 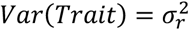 represents the stable characteristics of the subject, which may affect the ERP outcome; 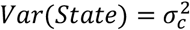 represents the subject’s psychological state, which may affect the ERP, and 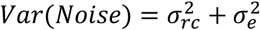 represents the residual error. Partitioning variance into these components was applied in a pointwise way along with the spatial and temporal domains of the ERPs.

#### 2.2.3 Statistical analysis

To further investigate the group- and individual-level measures in the reliability analysis, we analyzed the statistical consistency of reliability with the absolute value of the t-value and between-subject variance. Taking AEP as an example, one-sample t-test was performed along the post-stimulus time course at electrode Cz against a zero mean for grand average data on 82 subjects to derive group-level statistics. Time points that satisfied the Bonferroni-corrected criterion (p-value < 0.05/1000; 1000 was the number of post-stimulus time points) were selected to reduce the influences of noisy background activity. At these selected time points, the group- and individual-level measures were extracted, which were the absolute value of the t-value and standard deviation across the 82 subjects. Then the linear trends were removed from the time series of each measure and reliability to avoid spurious correlation. The associations between different measures and reliabilities were quantified by Spearman’s rank correlation coefficient, which is more robust to the non-linearity of changes and outliers than Pearson’s correlation. For SEP, VEP, and P300, same procedures were applied at electrodes Cz, Oz, and Pz, respectively, to explore the consistency between group effects and individual reliability because those electrodes showing the strongest group-level response.

### 2.3 Model Simulation

In an ERP experiment, the brain can be treated as a black-box system. With a certain type of stimulus input, the output of the system can be measured by ERP recording. However, changes in the internal variables in the brain cannot be accurately detected. As a supplement to the real EEG data analysis, a dynamic model simulation allows us to further understand the internal mechanism of the brain. In this work, a simulation model was applied to explore underlying factors that have a critical impact on the test-retest reliability of ERP analysis. David et al. (2005) considered that evoked changes of an EEG signal could be ascribed to transients that arise as the system’s trajectory returns to its attractor, which was more like a dampened oscillation and may not be necessarily associated with system parameter change. Considering that rhythmic oscillations are the basic characteristics of an EEG signal, a two-dimensional linear dynamical model, as shown in Eq. (4), was proposed in this work for the simulation of ERP. The linear dynamical model is:

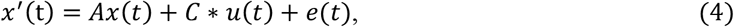

in which 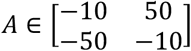 is the state-transition matrix with the corresponding eigenvalues −10 ± 50*i*.

The real parts of the eigenvalues were negative to ensure that the system would eventually converge to its point attractor [0,0], and the imaginary parts of eigenvalues indicated the periodical oscillation during the convergence. The input strength, *C*, was formulated as *C_sub_* + *C_trial_*, where *C_sub_* is a random variable representing the input strength for a given subject conformed to a Gaussian distribution (μ_sub_,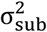), and *C_trial_* is a random variable representing the input strength for a given trial conformed to a Gaussian distribution (μ_trial_, 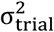). According to Jansen and Rit’s neural mass model (Jansen and Rit, 1995), the input of the system was simulated by using Eq. (5):

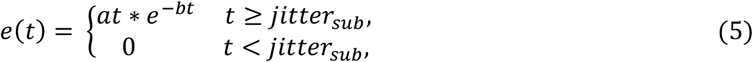

in which *jitter_sub_* is a rounded random variable with a uniform distribution [−*τ_sub_*, *τ_sub_*] relative to the onset time *t* = 0, and *e*(*t*) is the pink Gaussian noise representing the input of the background EEG activity in the simulation. The core setting of this model was the additive term *C_sub_* + *C_trial_*, which coupled the input strength with the subject-level and the trial-level, thus allowing both *Var*(*Trait*) and *Var*(*Noise*) to co-vary with the signal amplitude.

To better simulate the real data, there were 82 subjects with 120 trials for two sessions in the simulation. For the two sessions, there were 60 trials for each session. No systematic state variance was introduced considering the neglectable proportion of *Var*(*State*) in real data results. To mimic the real data preprocessing procedure, baseline correction was also applied to simulated ERP. With a sampling rate of 1000 Hz, there were 1500 time points for each trial, from −0.5 to 1 s. In this work, two major parameters of this model potentially influencing the test-retest reliability were investigated: (1) inter-subject variability, *τ_sub_*, for the latency jitter, *jitter_sub_*, and (2) inter-trial variability, *σ_trial_*, for the input strength, *C_trial_*.

Considering the state equation Eq. (4) with the phase portrait shown in Fig. 2(C), two factors, *τ_sub_* and *σ_trial_*, had a consistent form. Controlled by *σ_trial_*, the disturbance of input intensity would cause a change in the amplitude of the ERP response. Hence, it could be treated as a disturbance to the trajectory of the response from normal direction. While controlled by *τ_sub_*, the disturbance in the time domain did not change the amplitude of the ERP response but shifted its occurrence time in the phase portrait. Hence, it could be treated as a disturbance to the trajectory of the response from a tangential direction. Two factors, *τ_sub_* and *σ_trial_*, were selected to investigate the test-retest reliability in the simulation because they provided disturbances perpendicular to each other, in which their influence in different phases of the ERP response would be different. Further, *σ_trial_* is a trial-level factor, *τ_sub_* is a subject-level factor, and the change in *Var*(*Trait*) and *Var*(*Noise*) could be investigated in the simulation.

**Fig. 2.**
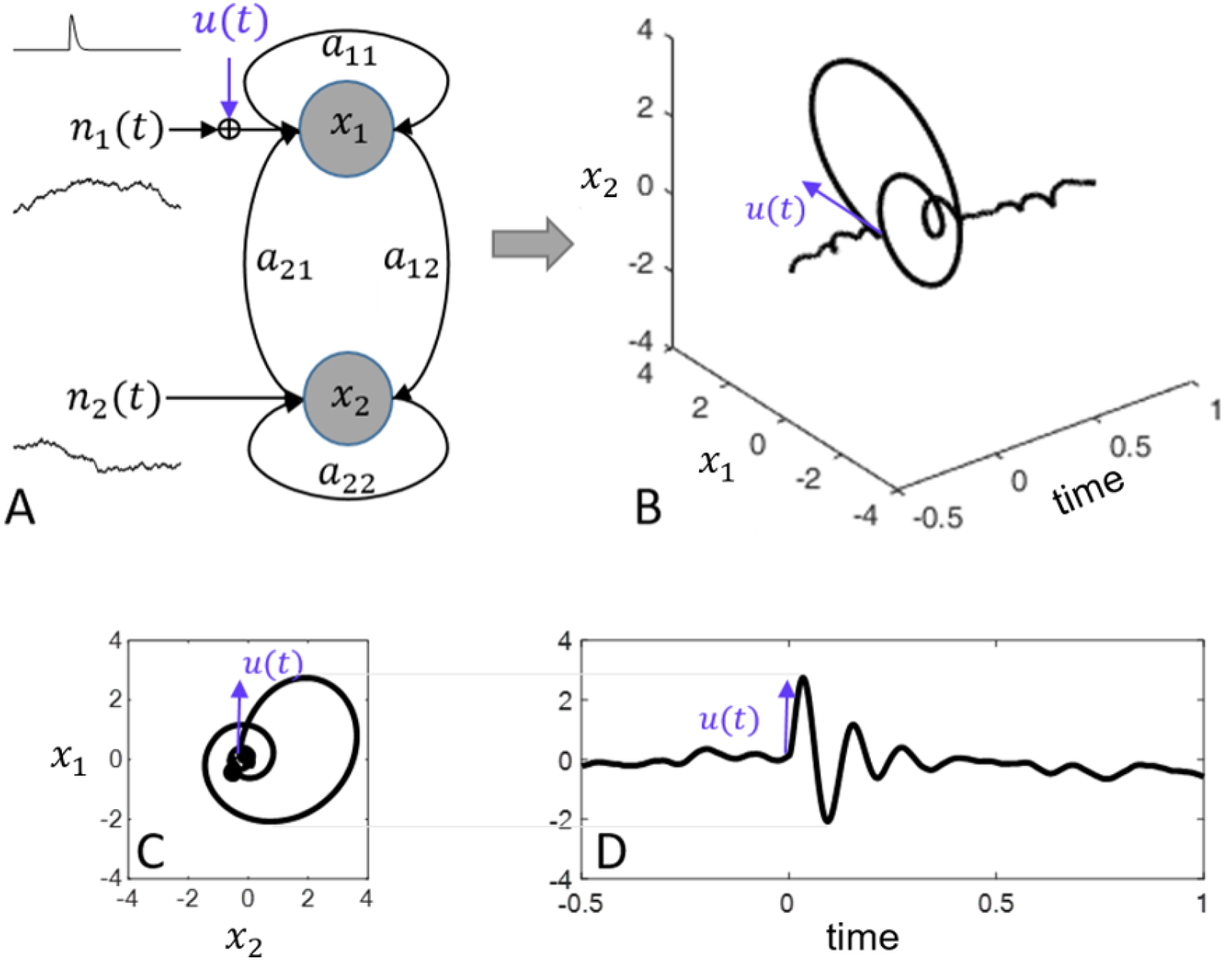
The ERP generation simulated by a second-order dynamic system model. (A) the framework of the dynamic model in Eq. (4); (B) the evolution of *x*_1_ and *x*_2_ over time, with its two-dimensional projection of the phase portrait in (C) and the evolution of *x*_2_ over time in (D).

## 3. Results

### 3.1 Reliability of real data

#### 3.1.1 Reliability for multisensory and cognitive ERPs

The grand average waveform of AEP at channel Cz, SEP at channel Cz, VEP at channel Oz, and P300 at channel Pz are shown in Fig. 3, where red and purple curves denote the signals of the two sessions. The representative ERP peaks included: N1 at 90 ms and P2 at 180 ms for AEP, N2 at 150 ms and P2 at 245 ms for SEP, N1 at 64 ms, P2 at 185 ms for VEP, and P3 at 345 ms of P300. The negative and positive peaks are respectively indicated by blue and yellow dashed lines in Fig. 3. For the pointwise analysis, the red dashed lines indicate the maximal reliability along the ERP time courses. As the subjects were more familiar with the experimental environment in the second session, the amplitudes of the ERP peaks were reduced.

**Fig. 3.**
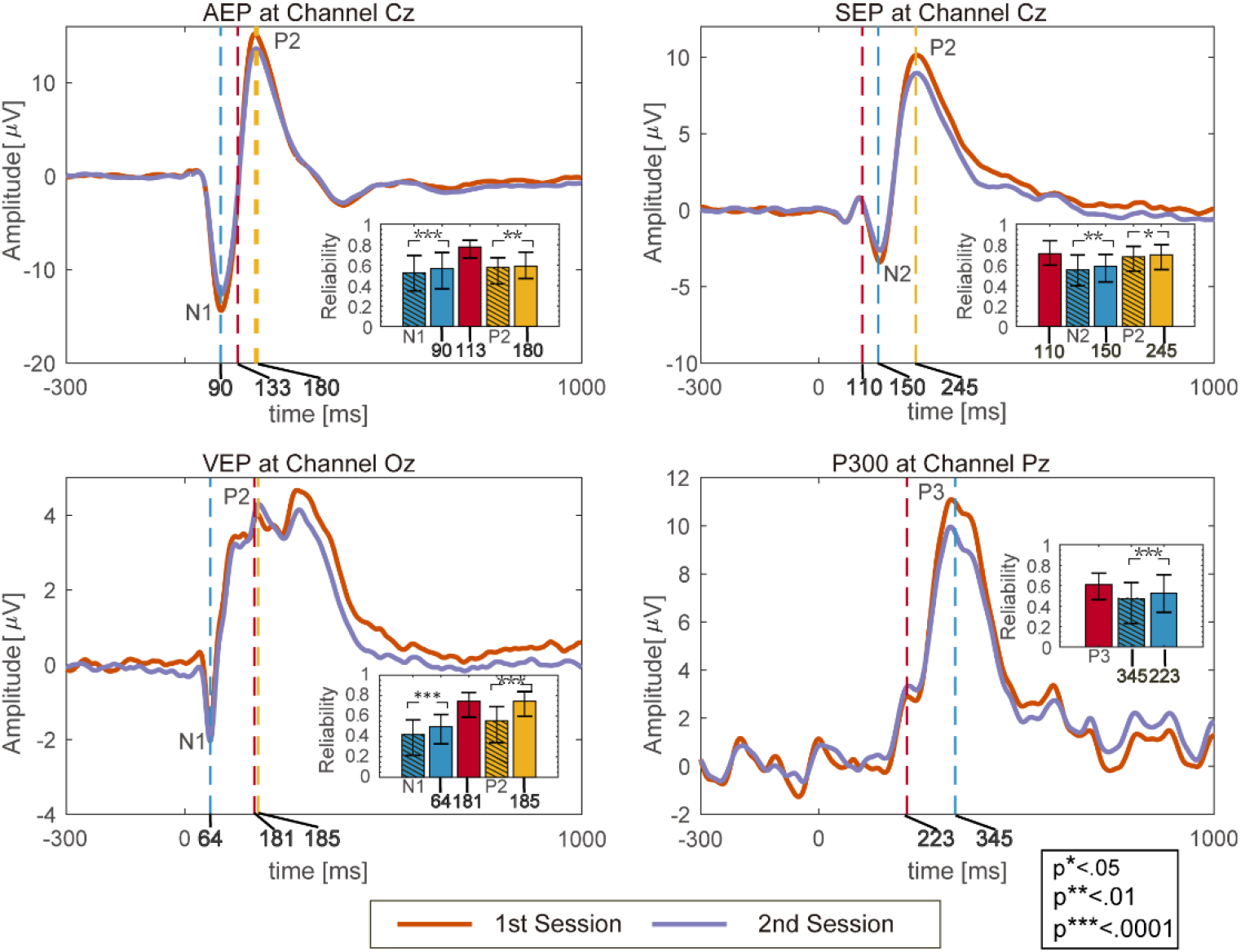
Grand average waveforms of the four types of ERPs at their representative channels: (A) AEP at channel Cz, (B) SEP at channel Cz, (C) VEP at channel Oz, and (D) P300 at channel Pz from the two sessions. The name of ERP (i.e., “N2”) in each bar plot represents the reliability of the peak amplitude of the ERP. The digit in bold font (such as “90”, “113”, and “180” in the subplot of AEP) in each bar plot represents the reliability obtained by extracting the amplitude of each subject at that time point as an individual difference variable.

As illustrated in the bar plots of Fig. 3, the reliability results of the peak amplitudes were compared with the pointwise analysis at corresponding time points. The results showed that the peak amplitude was significantly (p < 0.05) less reliable than the corresponding pointwise amplitudes at the latency of the grand average for all types of ERPs. Importantly, the most reliable time point in the ERP did not correspond to the ERP peak. For the VEP shown in Fig. 3C, the maximal reliability time point appeared at 182 ms, which was close to the peak of P2. For P300 shown in Fig. 3D, the maximal reliability time point in the pointwise analysis appeared at 220 ms (red dashed lines), with a reliability of 0.61 and bootstrap confidence interval of [0.46, 0.72]. This was much earlier than the well-known P3 component in the peak-based analysis, with a reliability of 0.47 and bootstrap confidence interval of [0.23, 0.63]. More interestingly, the time point with the maximal reliability for AEP appeared at 133 ms, with the mean amplitude being close to 0. The specific value in the bar plot can be found in Table.S1 in the Supplementary Material. Then, we used AEP as an example to conduct an in-depth analysis of the dynamic changes about the reliability of ERP, and similar results for VEP, SEP, and P300 can be found in the Supplementary Material.

#### 3.1.2 Spatiotemporal evaluation of reliability: a case study of AEP

Next, AEP was used to further investigate the consistency between group effects and reliability with different exploratory analyses (results for other types of ERPs are provided in the Supplementary Material). As illustrated in Fig. 4A, the t-value of significant regions that satisfied the Bonferroni-corrected criterion (p-value < 0.05/(1000*64, where 1000 is the number of post-stimulus time points, and 64 is the number of channels) are presented in the shaded region, which was consistent with the amplitude of the grand average waveform. Also, the post-stimulus AEP response behaved as a process of attenuating oscillations and finally approached the baseline. In contrast, the reliability of AEP after the stimulation shown in Fig. 4B increased significantly at first, lasting for a certain period, and then slowly returned to 0. Hence, the reliability of AEP along the ICC temporal profiles was not correlated with the amplitude of AEP. The maximal reliability of 0.78 appeared at the time point 133 ms, with the mean amplitude of AEP close to 0, which did not correspond to N1 at 90 ms or P2 at 180 ms with the minimal/maximal amplitude. The topographies of the grand average amplitude and the reliability at 90, 133, and 180 ms are illustrated in Figs. 4A and 4B.

**Fig. 4.**
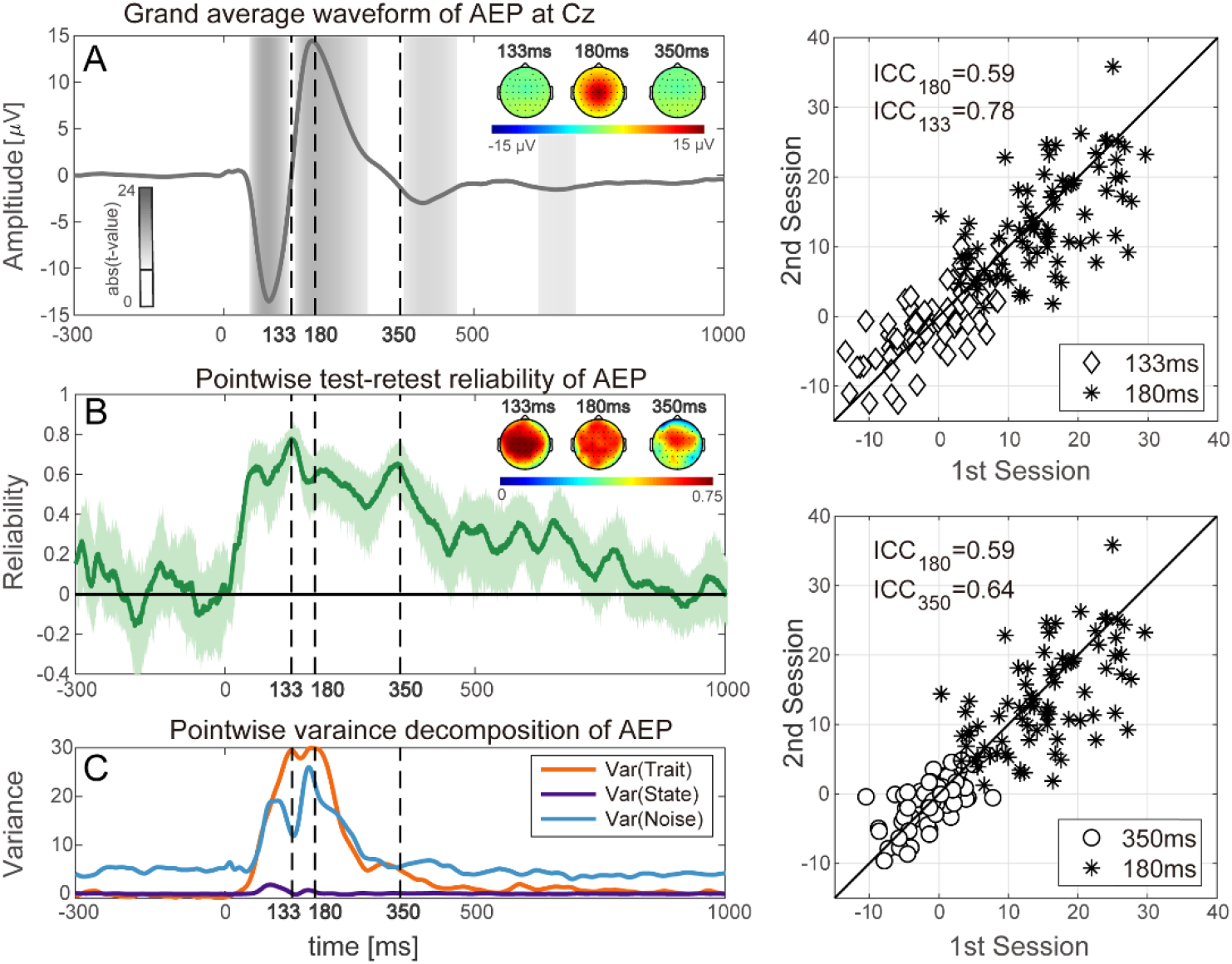
(A) Grand average waveform of AEP at electrode Cz. (B) Pointwise test-retest reliability analysis considering the entire shape of the AEP time-course calculated by ICC(A,1). (C) The variance in the observation matrix with the size of subject × experiment was decomposed into three parts: *Var*(*Trait*), *Var*(*State*), and *Var*(*Noise*) by a two-way random effects model along with the AEP time-course. (D) and (E) Scatter plots of 82 subjects’ amplitudes of AEP at electrode Cz in the two experiments were compared between 180 and 133 ms in (D) and between 180 and 350 ms in (E).

Pointwise variance decomposition results based on a two-way ANOVA are shown in Fig. 4C. The magnitude of *Var*(*Trait*) was close to 0 before the stimulation. At the beginning of stimulation, the magnitude of *Var*(*Trait*) reached a peak in the at 180 ms and then returned to 0. The local maximum of *Var*(*Trait*) did not correspond to the peak of AEP. The magnitude of *Var*(*Noise*) showed similar trends as *Var*(*Trait*), but the baseline was not 0. During the first 400 ms after stimulation, there was a certain correspondence between the waveform of AEP and *Var*(*Noise*). The peak of AEP corresponded to the local maximum of *Var*(*Noise*), while the zero-crossing point of AEP corresponded to the local minimum of *Var*(*Noise*). Compared with *Var*(*Noise*) and *Var*(*Trait*), the magnitude of *Var*(*State*) was too small and had little impact on reliability. Hence, the reliability was mainly determined by the ratio of *Var*(*Trait*) to *Var*(*Noise*). Next, the time points of 133, 180, and 350 ms were selected for the comparison, in which 180 ms corresponded to the peak of the grand average of AEP, while 133 and 350 ms corresponded to the local maximum of the reliability.

Figure 4D shows the comparison of the scatter plots of 82 subjects’ amplitude of AEP between time points 133 ms (diamonds) and 180 ms (asterisks). As *Var*(*State*) was close to 0, *Var*(*Trait*) could be measured as the variance along the black diagonal line, and *Var*(*Noise*) could be measured as the variance perpendicular to the black diagonal line. As shown in Fig. 4D, the mean amplitude of AEP at 133 ms (mean value of the diamonds) was much smaller than that at 180 ms (mean value of the asterisks), but the *Var*(*Trait*) values at the two different time points were similar. Hence, the reliability at 133 ms was larger than that at 180 ms because of the smaller *Var*(*Noise*) at 133 ms than that at 180 ms. Figure 4E shows a different situation compared with that shown in Fig. 4D. The reliabilities at 180 ms (asterisks) and 350 ms (circles) were similar, but both *Var*(*Trait*) and *Var*(*Noise*) at 350 ms were smaller than that at 180 ms.

#### 3.1.3 Statistical results

Spatiotemporal dissociation between group effects and individual reliability was revealed as shown in Figs. 3 and 4. These findings went against our expectations, given the fact that extracting peak-based measure using group-level prior information is the most common approach in reliability analysis. Hence, Spearman’s rank correlation analysis was further performed on AEP, SEP, VEP, and P300 to analyze the statistical relationships of reliability with the absolute value of the t-value and between-subject variance. In the results, there was a negative rank relationship between the the absolute value of the t-value and reliability concerning AEP with ρ = −0.14 and *p* = 0.01. For other ERPs, it still showed a weak relationship, as ρ = 0.26, *p* < 0.001 for SEP; ρ = 0.16, *p* < 0.001 for VEP; and ρ = 0.56, *p* < 0.001 for P300. Compared with group-level measure (t-value), the Spearman’s ρ between the individual-level measure (between-subject variance) and reliability was greatly improved.

### 3.2 Reliability of simulated data

#### 3.2.1 Simulation results

To further understand the internal factor influencing the reliability in ERP analysis, a dynamic model, as expressed by Eq. (4), was used for the simulation. The simulation results, shown in Fig. 5, were consistent with the results from real ERP data, shown in Fig. 4. Specifically, the grand average waveform of the simulated ERP is shown in Fig. 5A, with a peak of 134 ms and subsequent zero crossings appearing at 164 and 229 ms. The reliability curve across time is shown in Fig. 5B, and the corresponding variance decomposition is shown in Fig. 5C. In the simulation, the correlation between *Var*(*Noise*)and the amplitude of the ERP was more obvious. *Var*(*State*) was close to 0 because the systematic differences between the two sessions were not considered in this simulation. Hence, the reliability at the peak latency was the local minimum, and the reliability at the zero-crossing point was the local maximum. Similarly, the scatter plots in Figs. 5D and 5E show that the larger amplitude of the ERP may not necessarily lead to greater reliability, which was determined by the ratio of *Var*(*Trait*) to *Var*(*Noise*).

**Fig. 5.**
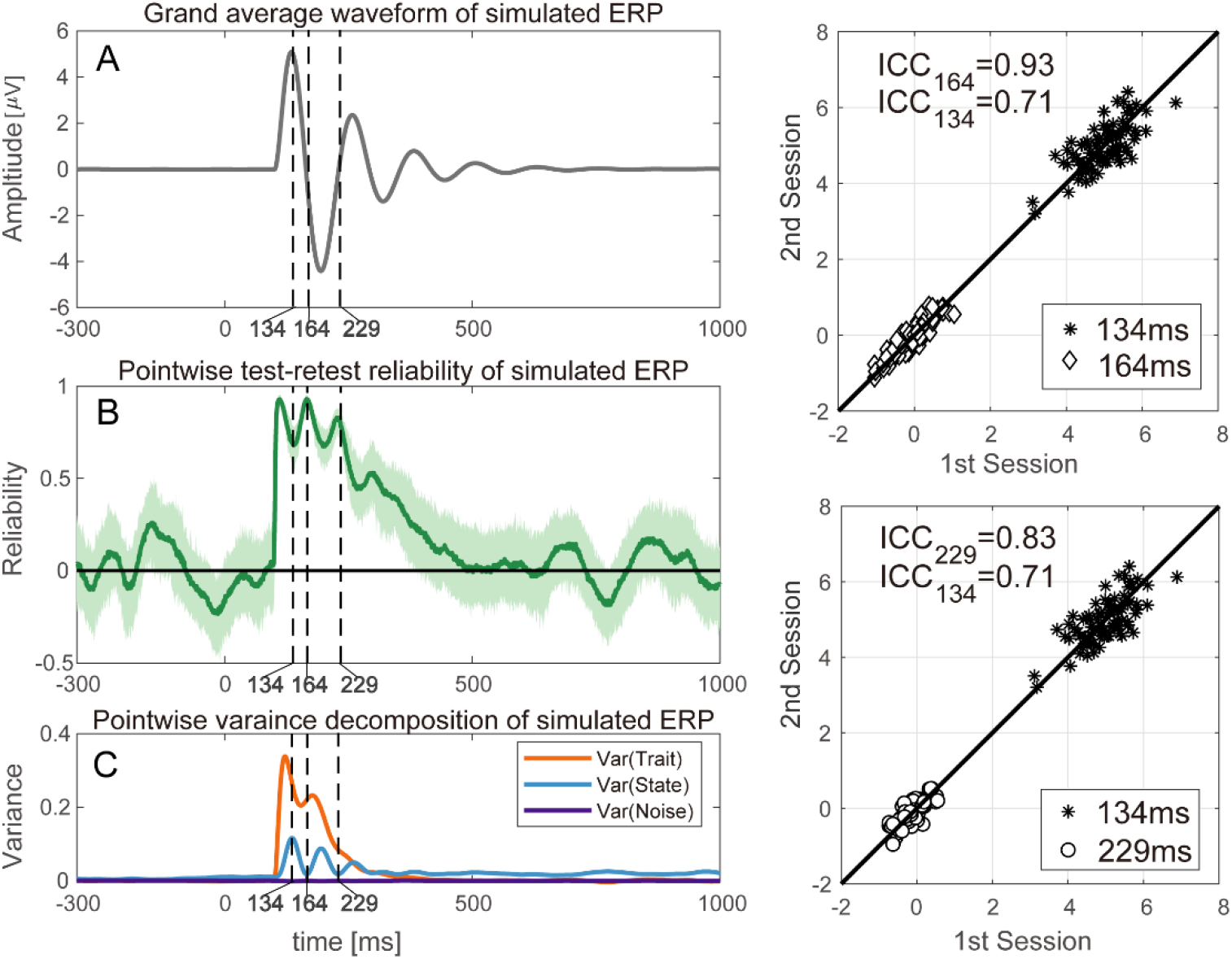
Grand average waveform of simulated ERP for a given set of system parameters (A). Pointwise test-retest reliability analysis along with the simulated ERP time course (B). The variance of observation matrix with the size of subject by experiment was decomposed into three parts: *Var*(*Trait*), *Var*(*State*), and *Var*(*Noise*) by a two-way random effects model along with the simulated ERP time-course (C). (D) and (E) Scatter plots of 82 subjects’ amplitudes of AEP at electrode Cz in two experiments were compared between 134 and 164 ms in (D) and between 134 and 229 ms in (E).

#### 3.2.2 The influence of the variability of jitter: *τ_sub_*

As a tangential disturbance in the phase portrait of Eq. (4) shown in Fig. 2, an increase in *τ_sub_* did not make a large difference in the waveform of the grand average ERP in the simulation, but it made the peak of P1 and N2 smoother. As *τ_sub_* increased from 0 to 20, the amplitude of the peak P1 reduced slightly, as shown in Fig. 6A. As illustrated in Fig. 6B, the reliability of the peaks at 134 and 194 ms remained around 0.7, while the reliability of the zero-crossing points at 164 and 229 ms increased greatly. As inter-subject latency jitter increased, the ICC values at 134 and 194 ms, which corresponded to the peaks of the grand average waveform, gradually shifted from the peaks of the ICC temporal profiles to their local minimum. The ICC values at 164 and 229 ms, which corresponded to the zero-crossing point, behaved conversely. The corresponding variance decomposition is shown with different values of *τ_sub_* in Fig. 6(D–E). *Var*(*Trait*) at the zero-crossing point (164 and 229 ms) of the grand average waveform increased as inter-subject latency jitter increased, while *Var*(*Noise*) fluctuated randomly. In comparison with the reliability of the peak amplitude, there was a greater difference between the maximum values of the ICC temporal profiles and the reliability of the peak amplitude.

**Fig. 6.**
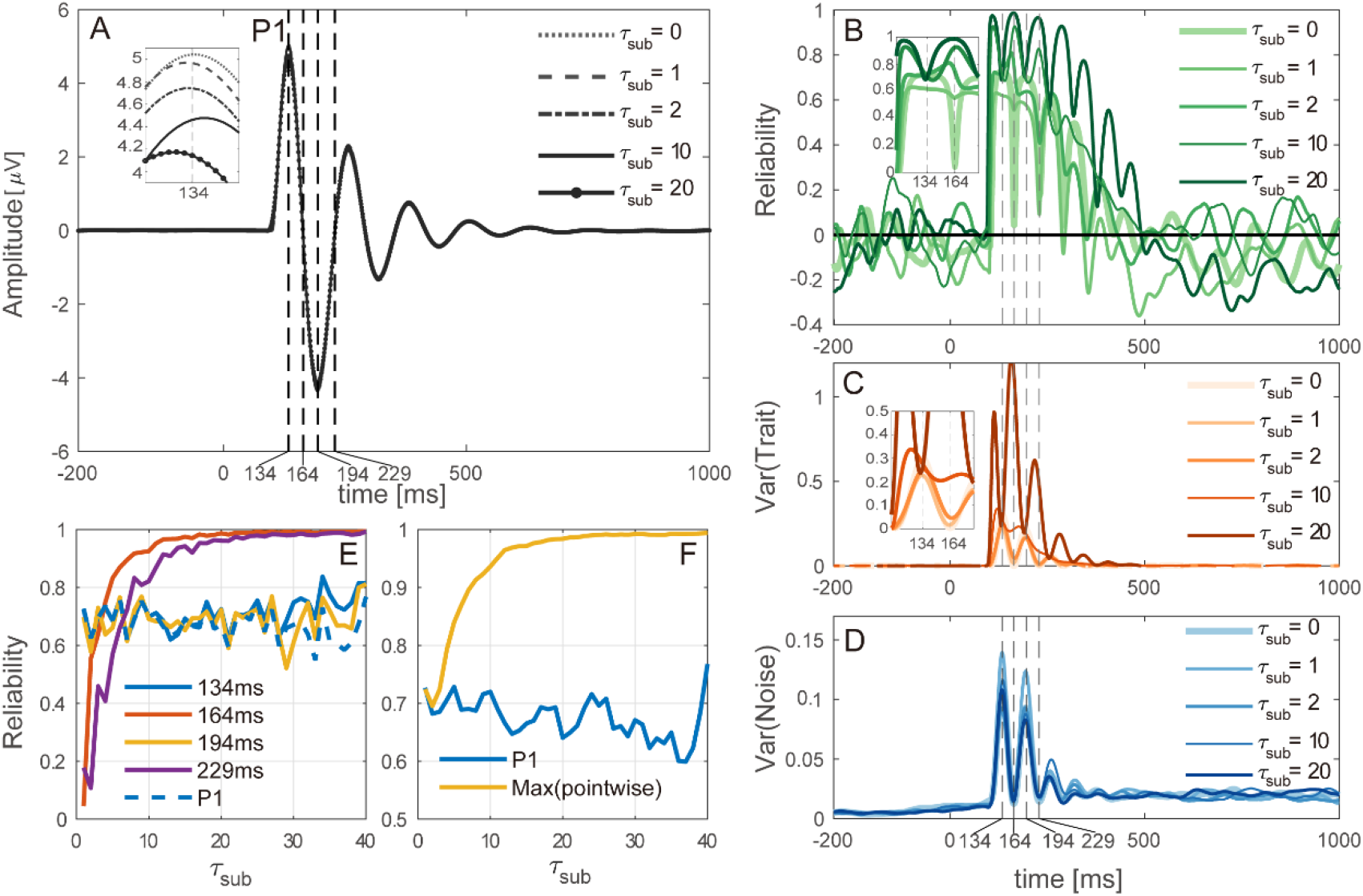
The influence of increasing the variability of inter-subject latency jitter of the dynamic system at the subject-level on (A) Grand average waveform of simulated ERP. (B) Pointwise test-retest reliability along the time-course of simulated ERP. (C) *Var*(*Trait*) along the time-course of simulated ERP. (D) *Var*(*Noise*) along the time-course of simulated ERP. (E) Comparisons between peak-based reliability and pointwise reliability at group-level peak latencies. (F) Comparisons between the maximum value of pointwise reliability and peak-based reliability.

#### 3.2.3 The influence of the variability of input power: *σ_trial_*

For normal perturbation in the phase portrait of Eq. (4) shown in Fig. 2, it is shown in Fig. 6B that the overall magnitude of the ICC temporal profiles dropped because of increasing inter-trial variability in the dynamic systems’ input, while the reliability in the response amplitude at 134 and 194 ms, which corresponded to the peak latencies of the grand average waveform, dropped more quickly compared with the reliability of the response amplitude at 164 and 229 ms, exhibiting an unbalanced influence. The above observations were further investigated by pointwise variance decomposition, in which the within-subject variation (*Var*(*Noise*)) increased systematically in proportion to the signal amplitude of the grand average waveform with increasing inter-trial variability, while the between-subject variation fluctuated randomly, thus explaining why the reliability of the signal with larger amplitude decreased more. Interestingly, it can be noted from Fig. 6F that there was a larger difference between the maximum values of the ICC temporal profiles and the reliability of the peak amplitude as the inter-trial variability increased.

## 4. Discussion

The purpose of this study was to investigate the relationships between group effects and individual reliability across different types of ERPs. By performing pointwise reliability analysis and rigorous simulation, we found inconsistency between individual reliability and group effects and provided potential explanations from the perspective of oscillations of ERP. The findings have implications for a series of questions that are of theoretical and practical relevance for ERP researchers, which will be discussed below sequentially.

### As the dominant approach in current ERP reliability studies, peak-based analysis has some potential problems

Until now, peak-related feature extraction (i.e., peak amplitude, area under the curve, mean amplitude) has been a dominant approach for examining the reliability of ERPs (Nordin *et al*., 2011; Munsters *et al*., 2019; Devos *et al*., 2020). For peak-based approach, researchers have found that the reliability of ERPs is influenced by the number of trials, channel selection, and various preprocessing strategies (Huffmeijer *et al*., 2014; Leue *et al*., 2013). The basic hypothesis behind the peak-based analysis is that the peak of the ERP indicates a higher signal-to-noise ratio, which produces results with a higher confidence level because of the relatively small interference from background EEG noise, yet this concept of signal-to-noise ratio may not generalize to the research area interested in individual difference, in which between-subject variance are treated as signal, within-subject variance are treated as noise, as mentioned by (Brandmaier et al., 2018). Another limitation of peak-based analysis is that the latency and amplitude of ERP peaks, as well as the entire ERP shapes, are physiologically meaningful and important (Gaspar *et al*., 2011).

### Spatiotemporal evaluation and decomposition of reliability are good for identifying reproducible individual difference

In this research, it was found that high signal-to-noise ratio assumption for the peak of ERP did not hold when considering individual difference research, which was also mentioned by Brandmaier *et al*. (2018). As illustrated in Fig. 4, the variance of the noise (blue curve in Fig. 4C) was highly correlated with the magnitude of the AEP response (absolute value of the black curve in Fig. 4A). Considering that the essence of EEG is neural oscillation, the peak in the ERP is just a certain phase (0 or *π*) during the oscillation. There is nothing more special about it compared to other phases. Hence, there is no reason why reliability analysis should be limited to peak-based features; pointwise analysis can bring us more comprehensive results. As compared with t-test or ANOVA, the pointwise analysis of test-retest reliability did not have the family-wise error rate problem, as we calculated the ICC values but did not judge whether there was a significant difference. Compared with peak-based analysis, the results from the pointwise analysis always had significantly higher ICC values at the time point of the peaks for all four types of ERP analysis in our investigation. In the test-retest reliability of AEP, SEP, VEP, and P300, the pointwise analysis consistently showed that the ICC value increased greatly after the stimulation, and after maintaining it for a while, decreased slowly to the baseline (Fig. 4). Hence, the peak of ERP may not relate to a higher ICC value. Even in AEP, the two local maximum points of the ICC value corresponded to the two zero-crossing points of the AEP. The findings suggest that reliability analysis restricted by the narrow time windows around the peaks is questionable. By performing pointwise analysis, dynamic changes in reliability in the spatial-temporal domain can be traced, given enough sample size, thus providing a new angle of ERP reliability analysis in a data-driven manner. Also, agreeing with the opinions of Gaspar *et al*. (2011), we believe that shape-based metrics rather than peak-based metrics may be more reliable for individual difference research.

### Stronger group effects do not guarantee higher individual reliability

In reliability analysis, the group effects are commonly used as a prior information (Plichta *et al*., 2012; Aron *et al*., 2006; Fliessbach *et al*., 2010), which assumes that experimental manipulation eliciting greater activation at the group level should also show reliable between-subject variation. This conventional approach has been questioned in recent years, especially in the fMRI community (Fröhner *et al*., 2019; Infantolino *et al., 2018;* Yarkoni and Braver, 2010; Li *et al*., 2019). In line with these studies, our results also revealed inconsistency between group effects and individual reliability in ERP analysis. More specifically, concerning the temporal domain discrepancies illustrated in Fig. 3, the most reliable points in the four types of ERPs (Cz for AEP, SEP, Oz for VEP, and Pz for P300) did not all correspond to maximum or minimum points of group-level activations. For AEP, the most reliable point appeared at the zero-crossing point of ERP. The spatial domain discrepancies are illustrated in Fig. 4 for AEP, in which we did not find the topography of the AEP response corresponding to the topography of the reliability at 133 and 350 ms. Further analysis, presented in Table 1, indicated that, as an individual-level measure, the between-subject variance showed a higher correlation coefficient than the group-level measure (abs(t-value)) across all four types of ERPs. All these evidences suggest that it is not advisable to select peak-related features at the electrode showing the strongest group effects without carefully examining their reliability.

**Table 1.**
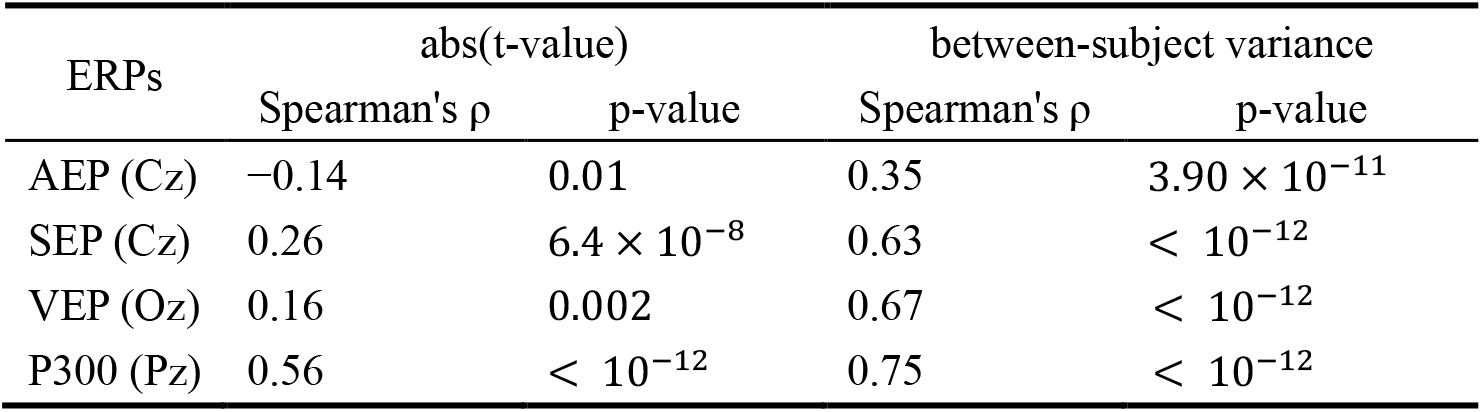
Associations between group-level measure (t-value), individual-level measure (between-subject variance), and reliability

| ERPs | abs(t-value) |  | between-subject variance |  |
| --- | --- | --- | --- | --- |
| | Spearman's $\rho$ | p-value | Spearman's $\rho$ | p-value |
| AEP (Cz) | -0.14 | 0.01 | 0.35 | $3.90 \times 10^{-11}$ |
| SEP (Cz) | 0.26 | $6.4 \times 10^{-8}$ | 0.63 | $< 10^{-12}$ |
| VEP (Oz) | 0.16 | 0.002 | 0.67 | $< 10^{-12}$ |
| P300 (Pz) | 0.56 | $< 10^{-12}$ | 0.75 | $< 10^{-12}$ |

Intuitively, the spatial-temporal distribution of group-level analyses and individual-level analyses should tend to converge. In other words, increased activation by experimental manipulation at the group level should relate to individual-level analysis, given a large enough sample size and no confounding factors. However, few empirical pieces of evidence support this idea (Lee *et al*., 2006); more often, individual difference analyses simply fail to reveal any significant effects in regions that show a robust within-subject effect (Vetter *et al*., 2017; Raemaekers *et al*., 2007). In this research, the simulation results indicate that the consistency between group-level effects and individual reliability may be dynamically modulated by inter-subject latency jitter and inter-trial variability of dynamic system input, providing a dynamic view of the relationships between the two types of analysis in ERP analysis.

### Both peak and zero-crossing points of ERPs just represent different phases of one unified oscillation process

To further understand spatial-temporal inconsistency between group-level effects and individual reliability in ERP analysis, a dynamic model was applied for the simulation. The simulation model was simplified to be a second-order linear attractor with noise to simulate the EEG oscillation. From the perspective of dynamic system theory (Jansen and Rit, 1995; Youssofzadeh *et al*., 2015), peaks in the EPR are just an observation of EPR from one dimension of the computational models of neural processes. The phase portrait of our simulations (Fig. 2) provides a more comprehensive perspective, in which the peaks are just some special phases during the neural oscillation. In the dynamic model, the variability of jitter *τ_sub_* provides a subject-level disturbance tangent to the trajectory of the ERP oscillation, and the variability of input power *σ_trial_* provides a trial-level disturbance perpendicular to the trajectory of the ERP oscillation. Owing to the different directions and different levels of the two factors, the simulation result showed that the changes of these two factors played very different roles in different phases of ERP. Figures 6 and 7 illustrate the effect, especially at the peak and zero-crossing points of the ERP, with the changes of these two factors. With the similar wave form of the group-level ERP, the reliability would be determined by several factors. Measuring the peak-based features would not provide a comprehensive understanding about the oscillations in ERP.

**Fig. 7.**
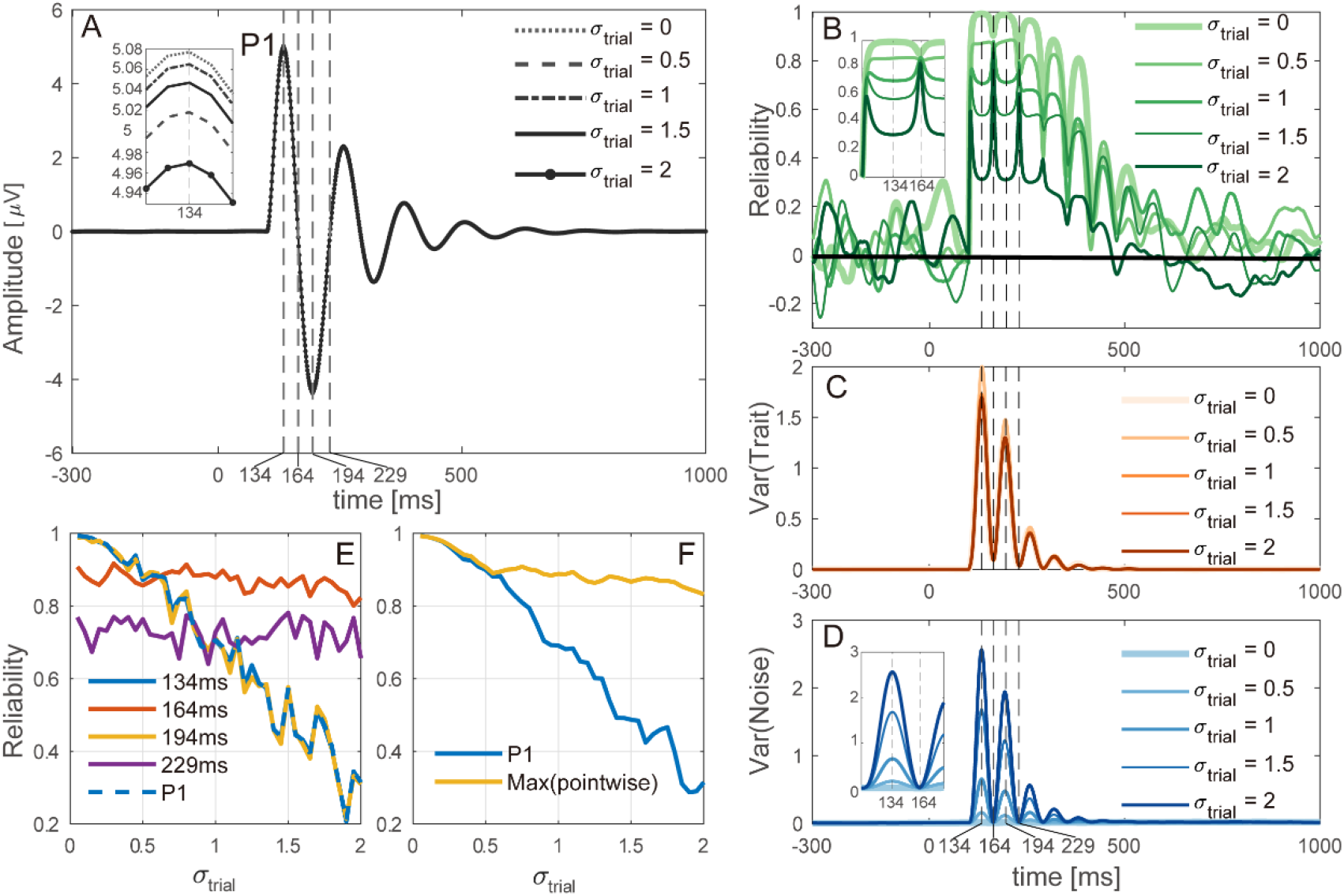
The influence of increasing the variability of input power of the dynamic system at the trial-level on (A) Grand average waveform of simulated ERP. (B) Pointwise test-retest reliability along the time-course of simulated ERP. (C) *Var*(*Trait*) along the time-course of simulated ERP. (D) *Var*(*Noise*) along the time-course of simulated ERP. (E) Comparisons between peak-based reliability and pointwise reliability at group-level peak latencies. (F) Comparisons between the maximum value of pointwise reliability and peak-based reliability.

### To translate group effects into individual difference research, some issues must be reconsidered in ERP data analysis

ERP analysis focusing on individual difference often implicitly or explicitly uses prior information from group effects. For reliability analysis, electrodes showing the strongest stimulus-related activity by group-level analysis are often chosen for test-retest reliability analysis of ERPs (Gaspar et al., 2011). For constructing single-subject predictive models, it has been done in translating findings in group-level statistics of ERPs into a machine learning framework (Boshra et al., 2019). Hence, here we discuss some critical issues about ERP data analysis concerning the use of group effects in individual difference research.

- Tracing back to history, the original idea of ERPs was to index specific cognitive processes rather than distinguishing different individuals (i.e., the research interest was how brain activity responds to one condition versus the other). Individual differences were treated as measurement error that could not be explained by experimental manipulation, as t-test, omnibus ANOVA assumes. From this perspective, there is no reason to select regions based on strongest group effects and then feed them into the correlation or reliability analysis, except that this region also shows greater between-subject variation.
- For the data analysis pipeline of ERPs, it is very common to perform the subtraction operation (e.g. ERP difference waves) to minimize the impact of baseline individual differences. Such operations forcibly promote the activity of the baseline period at a constant rate across individuals, and the goal is to obtain a reliable experimental effect at the group level. This pervasive practice was inherited from research focusing on experimental effects, but few studies have noticed whether this approach is reasonable for individual difference analysis. Recently, the fMRI and psychology communities have argued that difference scores often exhibit a robust group-level effect but lower reliability (Infantolino *et al*., 2018; Onie and Most, 2017). Considering statistical analysis, typically, ERP data are averaged within conditions and participants after preprocessing and then analyzed for the mean difference between conditions using paired t-test or repeated measures ANOVA. This traditional approach implicitly assumes that experimental manipulation yields uniform effects across all participants. Random variance of individual difference in effect sizes is not taken into account. By adopting linear mixed-effect models, in which random effects are used to capture individual variability as a form of random slopes or random intercepts, fixed effects are estimated by the grand mean across all participants. Such an approach has been adopted to simultaneously capture both group effects and individual difference (Frömer *et al*., 2018; Tibon and Levy, 2015).

### Several limitations are presented below

Our research and reliability analyses had several limitations. First, higher reliability does not ensure higher validity. The fact that the response amplitude at some time points was more reliable than the peak amplitude may be explained by sacrificing validity. More specifically, each subject’s response amplitude at a given time point may indexed different neurophysiological processes, leading to larger between-subject variance. Increasing reliability in this way is not desirable because the underlying process of this measure is different across subjects. However, we cannot verify this potential explanation without behavioral data. Second, the neglectable portion of *Var*(*State*) may be attributed to not having enough sessions to capture systematic variance. Third, our analysis was restricted to univariate features; the relationship between group-level effects and individual reliability concerning multivariate analysis warrants further investigation in the future.

## 5. Conclusion

In summary, the purpose of this research was to investigate the consistency between group effects and individual reliability of ERPs. We performed spatiotemporal evaluation and decomposition of reliability in four different ERPs, and the findings indicate that peak-based approach (i.e., selecting regions showing the strongest group-level response as individual difference variables) may be inappropriate for reliability analysis of ERPs. Without carefully examining reliability, this approach based on group-level prior information may fail to reliably capture individual differences, which is supported by spatiotemporal dissociation between group effects and individual reliability. The disadvantages of peak-based reliability analysis were illustrated by spatiotemporal evaluation and decomposition of reliability, statistical results and the phase portrait in the simulation model. Further, the simulation results highlight the modulation role of inter-subject latency jitter and inter-trial variability in modulating the consistency between group-level effects and individual reliability. To conclude, all these results provide a new perspective beyond peak-based analysis in the ERP reliability studies. Furthermore, the findings deepen our understanding of ERP generation and the reliability of ERPs.

## Acknowledgments

This work was supported by the National Natural Science Foundation of China (No. 81871443), the Science, Technology and Innovation Commission of Shenzhen Municipality Technology Fund (No. JCYJ20190808173819182), and Shenzhen Peacock Plan (No. KQTD2016053112051497).

None of the authors has potential conflicts of interest to be disclosed.

